# Loss of long-range co-expression is a common feature in cancer

**DOI:** 10.1101/2022.10.27.513947

**Authors:** Diana Garcia-Cortes, Enrique Hernández-Lemus, Jesús Espinal Enríquez

## Abstract

Cancer cells display common features and enabling characteristics collectively known as the Hallmarks of Cancer, which occur alongside alterations in the regulatory mechanisms controlling gene transcription. Gene co-expression networks (GCNs) provide a framework to identify correlated gene sets that may share these regulatory mechanisms. Previously, we reported the loss of long-range co-expression in breast, lung, kidney, and hematopoietic cancer GCNs. Here, we expand the analysis to fifteen tissues, comprising 8,772 samples from two independent datasets. Unlike healthy phenotypes, cancer GCNs show that the strongest gene-pair interactions are intra-chromosomal, with their strength decaying as base-pair distance increases. Tumor GCN communities are strongly associated with cancerrelated processes and are enriched in gene families located on the same chromosome. In contrast, normal GCN communities are linked to metabolic and cell maintenance processes. Riboproteins remain highly co-expressed in both cancer and normal GCNs, highlighting their importance for cell viability. Notably, in other chronic diseases, such as Type-2 Diabetes and Alzheimer’s disease, the loss of long-range co-expression is absent, suggesting it is a distinctive feature of cancer.

## Introduction

The Hallmarks of Cancer (1), a conceptual framework that has evolved over the past two decades, describe the distinctive characteristics that cancer cells acquire as they progress toward tumorigenesis (2–4). The authors also identified enabling characteristics, highlighting the role of genome instability, tumor microenvironment, and epigenetic alterations in facilitating the acquisition of hallmark capabilities (5, 6).

Alterations in multiple regulatory mechanisms drive the manifestation of cancer hallmarks. In gene transcription, these mechanisms include cis-regulatory elements, specialized proteins (e.g., transcription factors), and epigenetic markers, all operating within a specific chromatin configuration (7–9). Analyzing gene co-expression profiles can provide insights into altered regulatory mechanisms that affect the coordinated expression of multiple genes (10–12).

Gene co-expression profiles are generated by calculating the correlation between pairs of genes in a gene expression matrix derived from multiple samples of a given phenotype. The most significant pairs can be extracted to construct a gene coexpression network (GCN). GCNs provide a topological description of a transcriptome, enabling comparisons of global and local connectivity patterns across different phenotypes, as well as their functional implications (13–15).

By analyzing gene co-expression profiles and building GCNs using RNA-Seq data in cancer tissue and its normal counterpart, we have previously described a loss in the proportion of inter-chromosomal gene co-expression interactions in breast cancer and breast cancer molecular subtypes (16– 19), lung cancer (20), clear cell renal carcinoma (21), and hematopoietic cancers (22). There, the majority of significant co-expression links connect gene pairs in the same chromosome and these display a highly localized co-expression pattern. Conversely, their normal tissue-derived networks, present a higher number of inter-chromosomal interactions. Local high co-expression is a characteristic that has been reported for normal tissues, both in bulk RNA-Seq (23) and single cell analysis (24), reinforcing the conclusion that gene order in eukaryotes is not random (25, 26). However, this phenomenon has been observed in gene pairs within a local vicinity of less than 1Mb (27, 28). Our previous results indicate that in cancer, this high co-expression expands to longer distances and its prevalence throughout different tissues suggests it may be a common characteristic of the disease. Moreover, in terms of the regulatory mechanisms contributing to the emergence of high co-expression regions, we did not find a decisive association with a single mechanism in the Luminal A breast cancer subtype (29). Indeed, in normal tissues, multiple factors contributing to local co-expression have been reported (23). These results, in both cancer and normal phenotypes, highlight the need to extend the analysis of gene co-expression and the mechanisms that jointly drive the transcription of multiple sets of genes.

In this work, we simultaneously evaluated the loss of longrange co-expression in fifteen tissues (8,772 samples) by integrating data from distinct data sources: TCGA (30) and UCSC Xena (31) (which includes samples from the GTEx and TCGA collaborations (32)). This in order to determine whether (i) the loss of inter-chromosomal co-expression is a common feature in cancer, (ii) there is a dependency between high co-expression and physical distance between genes located in the same chromosome, and (iii) whether these phenomena are exclusively found in cancer. To do this, we analyzed gene co-expression profiles and GCNs from said samples, compared their topological properties through network community extraction, and identified biologically associated communities. This analysis allowed us to confirm that loss of long-range co-expression is indeed a common and distinctive feature in cancer.

## Results

### In cancer, highly co-expressed gene pairs are intra-chromosomal

We calculated Mutual Information (MI) between pairs of genes as a measure of co-expression for each tissue and sample type independently. The number of gene pairs analyzed varied by tissue, ranging from 49.07 million in skin to 108.86 million in lung, due to tissue-specific preprocessing filters. Table 1 displays the number of samples and genes in each expression matrix.

**Table 1.**
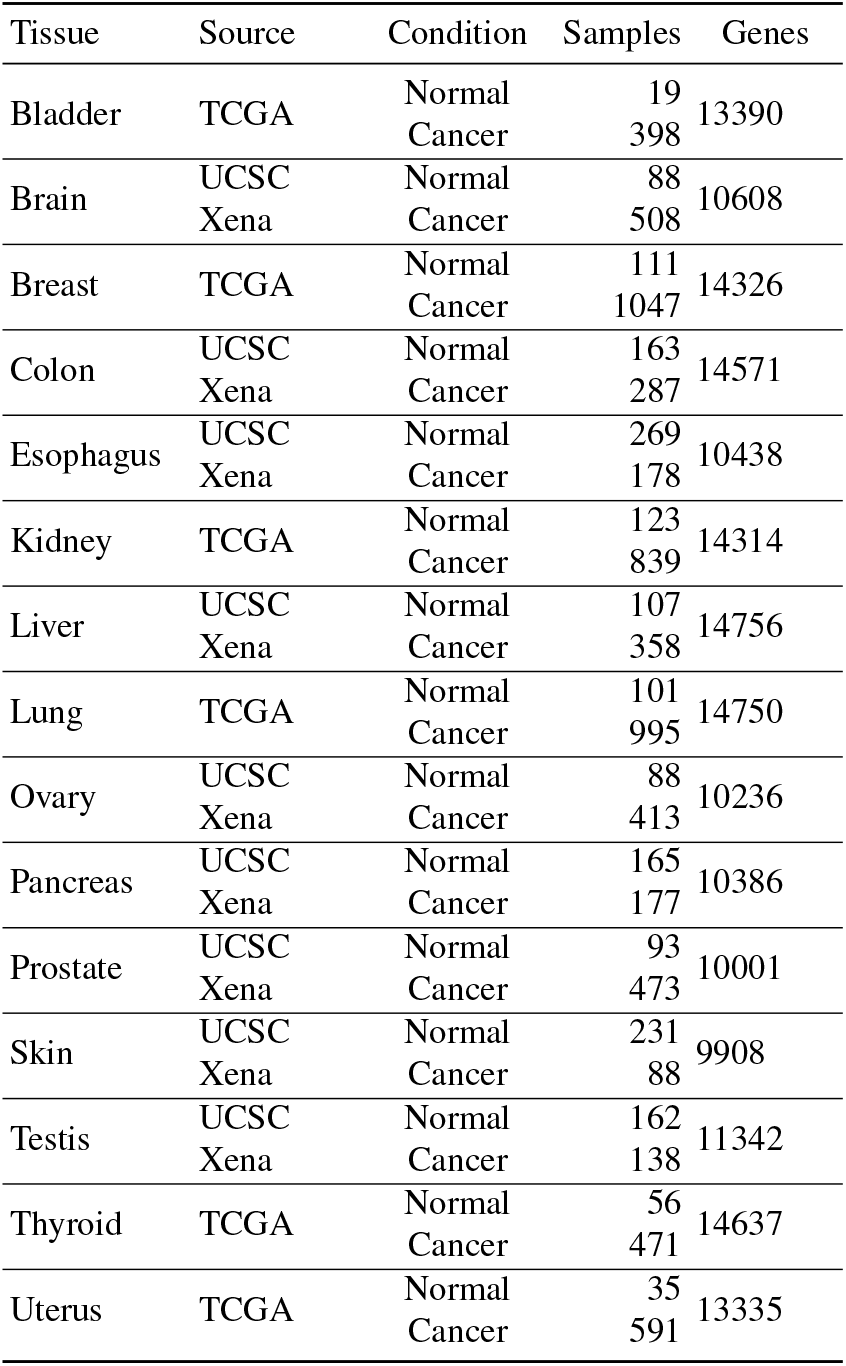
Number of samples and their data sources for each tissue in the study. The number of genes (Genes) in the expression matrix changes per tissue due to processing filters.

On average, 5.33% of gene pairs consist of two genes located on the same chromosome, representing intra-chromosomal interactions. In normal tissues, this percentage remains stable across different thresholds of top MI values. However, in cancer tissues, higher fractions of intra-chromosomal interactions appear among the top MI values. Figure 1 presents a heatmap showing the fraction of intra-chromosomal interactions for various thresholds of top MI values in both cancer and normal tissues. For bladder samples, these values are also depicted in a line plot, while Supplementary Figure 1, provides line plots for the remaining tissues.

**Fig. 1.**
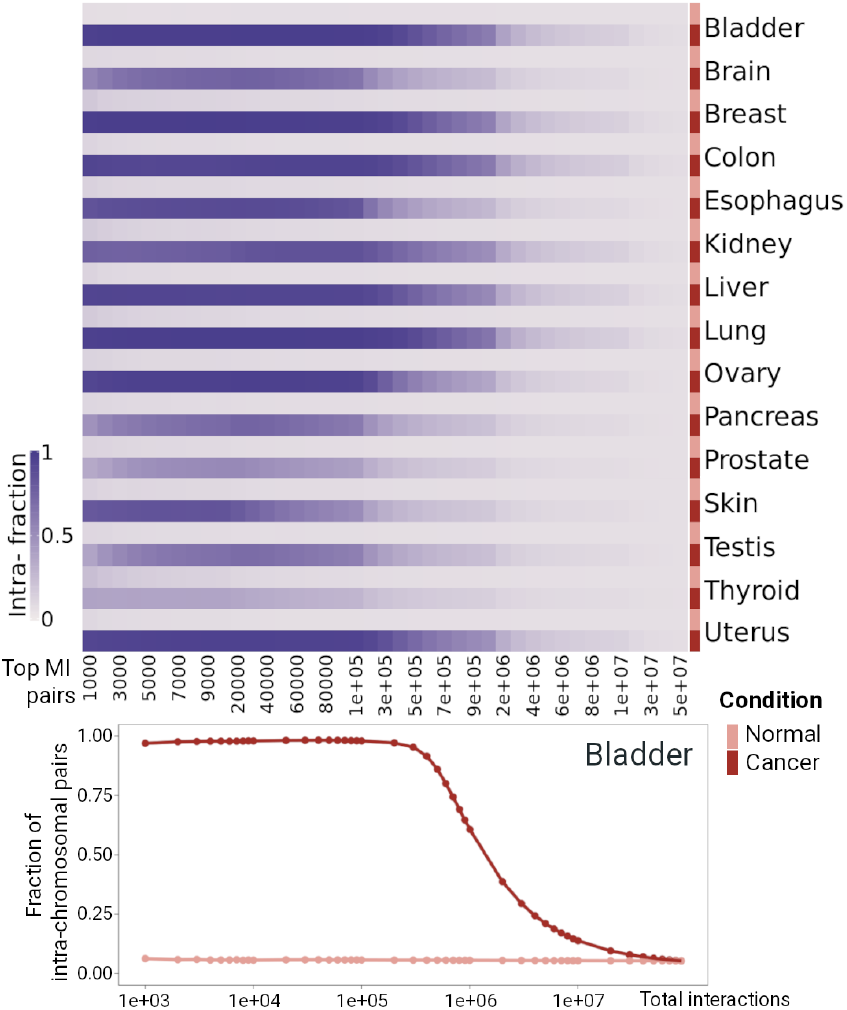
Heatmap showing the fractions of intra-chromosomal interactions at different top mutual information (MI) thresholds for cancer and normal tissues across all fifteen datasets. In normal co-expression profiles, this fraction remains stable across thresholds, whereas in cancer, the top MI interactions are predominantly intra-chromosomal. For bladder, the fractions are also displayed as a line plot, with the remaining tissues shown in Supplementary Figure 2.

Kolmogorov-Smirnov tests reveal significant differences in the distribution of intra-chromosomal fractions between cancerous and normal tissues, with p-values ranging from 1e-16 to 0.005 (Supplementary File 1) when considering the top 1 million interactions. This finding suggests that in all analyzed cancer types, the strongest co-expressed gene pairs are located on the same chromosome, whereas normal tissues do not show a preference for high co-expression among intrachromosomal interactions.

### Distance dependent MI decay is a common feature in cancer

After establishing that the strongest co-expression interactions in cancer occur between genes on the same chromosome, we investigated whether correlation values depend on the chromosomal distance between these intrachromosomal genes. We assessed intra-chromosomal coexpression by sorting gene pair interactions based on their base pair distance. Figure 2 displays the average MI values for 1,000-interaction bins, with corresponding standard deviations plotted against distance, in colon tissue. Supplementary Figure 2 provides similar plots for all fifteen tissues in the study.

**Fig. 2.**
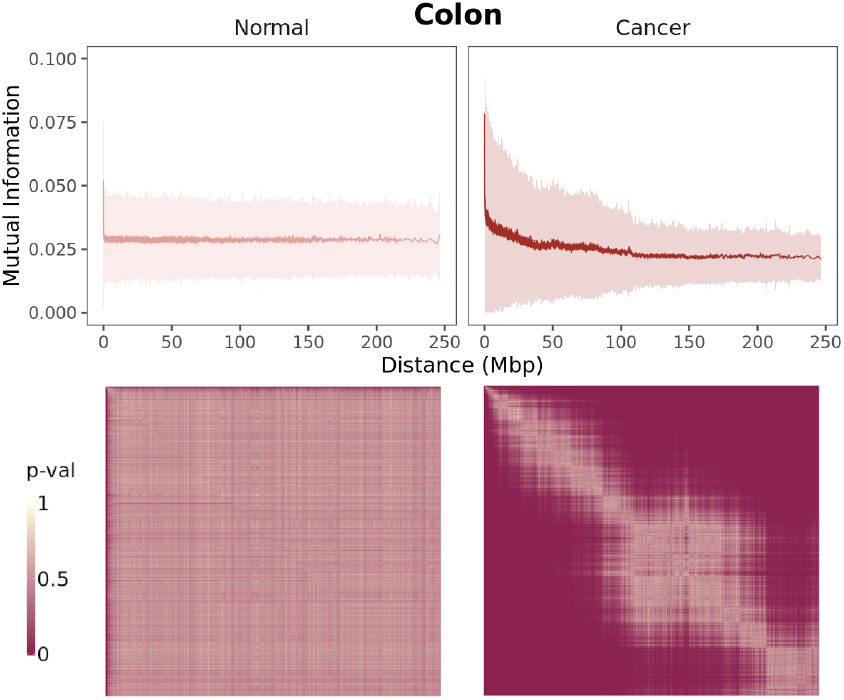
Average Mutual Information (MI) and standard deviation for bins of 1,000 intra-chromosomal interactions plotted against average gene-gene base pair distance in colon tissue. The lower panel shows p-values from Wilcoxon rank sum tests comparing MI distributions between bins. In normal tissue, no significant differences are observed between bins, while in cancer, distant bins differ significantly, but neighboring bins show similar values. Supplementary Figure 3 includes similar plots for the other tissues.

In cancer, gene pairs with shorter distances exhibit higher MI values, and as distance increases, co-expression decays to a plateau. In contrast, normal intra-chromosomal coexpression profiles show this pattern only for genes in very close proximity. We performed Wilcoxon rank sum tests to compare the distribution of MI values among bins. P-values, shown for colon cancer in the lower panel of Figure 2 and in Supplementary Figure 3 for the full cohort, indicate that in cancer, neighboring bins are similar, while distant bins differ significantly. Conversely, normal-derived bin distributions do not show differences related to base pair distance. Thus, not only do the strongest co-expressed genes reside on the same chromosome, but they are also placed in close proximity.

### Distinct intra-chromosomal interaction patterns in cancer co-expression networks

We used the top 100,000 MI interactions from normal and cancer tissues to construct gene co-expression networks (GCNs). Table 2 presents their main properties, while Figure 3 provides a visual representation of networks for selected tissues. Supplementary Figure 4 includes networks for all tissues. As previously noted, at this cutoff, most interactions in cancer networks involve gene pairs on the same chromosome, whereas normal networks resemble what we observed earlier in breast and lung cancer: a giant component primarily composed of inter-chromosomal links (16, 18, 20).

**Fig. 3.**
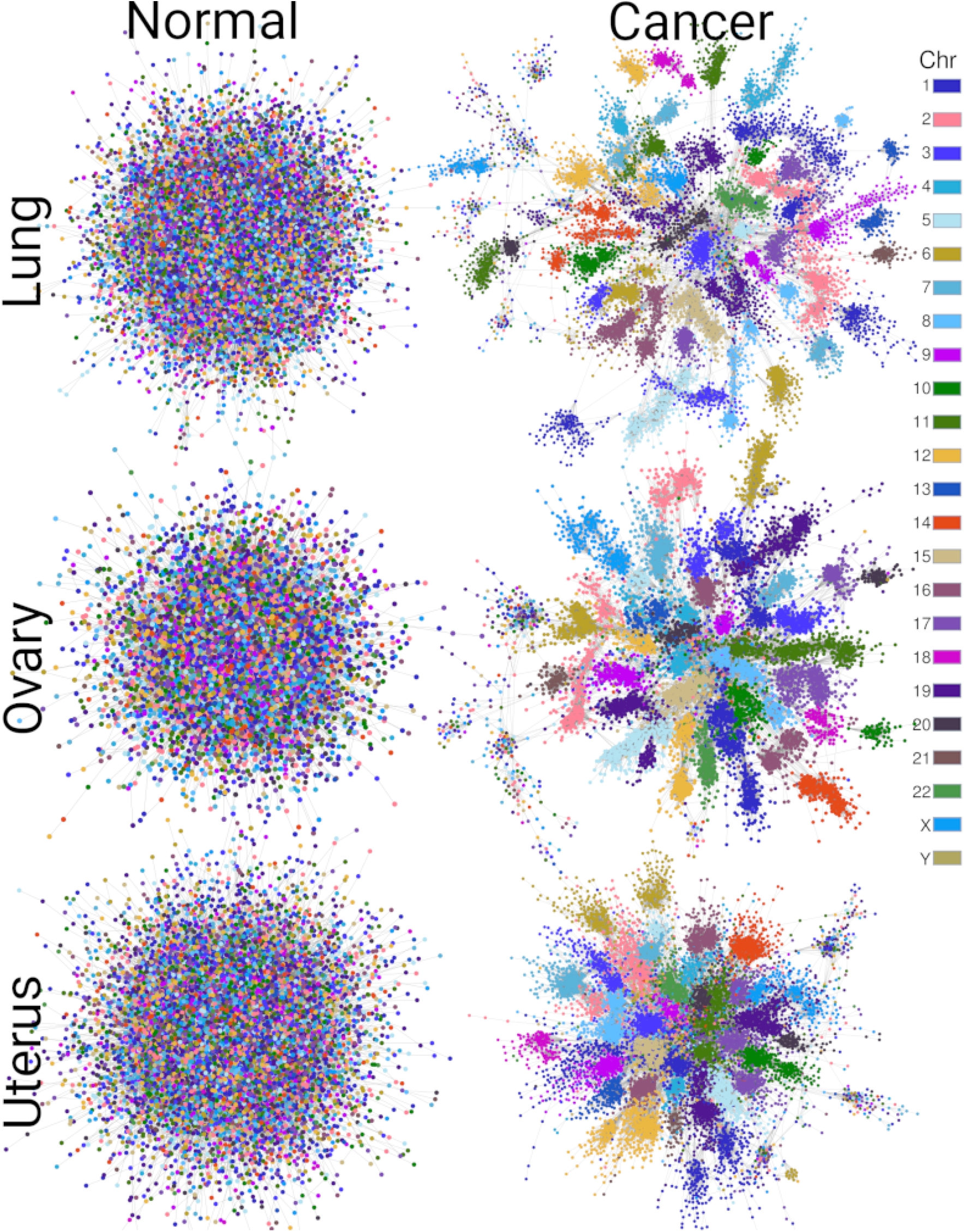
Co-expression networks from the top 100,000 interactions in selected tissues, visualized using a force-directed layout. Cancer networks exhibit modules dominated by intra-chromosomal interactions, a pattern not observed in normal networks. Supplementary Figure 4 shows networks for all tissues.

**Table 2.** Main features of normal and cancer co-expression networks for all fifteen tissues in the study. Networks are built by taking the top 100,000 MI interactions, although they differ in the number of nodes (Genes). The Intrafraction refers to the fraction of links joining genes in the same chromosome. Communities were computed using the Louvain algorithm (see). Up and Down refers to genes upregulated and downregulated (*pval*_*adj*_ < 0.05) in the differential gene expression analysis.

| Tissue | Condition | Genes | Intra- fraction | Communities | Modularity | Up | Down |
| --- | --- | --- | --- | --- | --- | --- | --- |
| Bladder | Normal | 13221 | 0.05709 | 34 | 0.554 |  |  |
|  | Cancer | 9403 | 0.98007 | 321 | 0.967 | 4618 | 1995 |
| Brain | Normal | 8897 | 0.06249 | 68 | 0.505 |  |  |
|  | Cancer | 8738 | 0.65396 | 129 | 0.796 | 4735 | 3436 |
| Breast | Normal | 11902 | 0.08626 | 93 | 0.515 |  |  |
|  | Cancer | 10155 | 0.97363 | 423 | 0.965 | 5571 | 3435 |
| Colon | Normal | 10645 | 0.07128 | 176 | 0.717 |  |  |
|  | Cancer | 8958 | 0.87639 | 373 | 0.923 | 5111 | 3264 |
| Esophagus | Normal | 8325 | 0.08036 | 85 | 0.624 |  |  |
|  | Cancer | 9117 | 0.83717 | 139 | 0.896 | 4580 | 3856 |
| Kidney | Normal | 10955 | 0.08052 | 89 | 0.556 |  |  |
|  | Cancer | 9612 | 0.85921 | 394 | 0.894 | 5168 | 3486 |
| Liver | Normal | 11952 | 0.07664 | 144 | 0.596 |  |  |
|  | Cancer | 9622 | 0.91591 | 357 | 0.945 | 4984 | 3551 |
| Lung | Normal | 12469 | 0.07914 | 80 | 0.545 |  |  |
|  | Cancer | 10116 | 0.97844 | 424 | 0.971 | 6950 | 2321 |
| Ovary | Normal | 9071 | 0.07302 | 44 | 0.448 |  |  |
|  | Cancer | 7729 | 0.95778 | 193 | 0.959 | 3626 | 3496 |
| Pancreas | Normal | 8057 | 0.06685 | 97 | 0.506 |  |  |
|  | Cancer | 9659 | 0.61381 | 51 | 0.746 | 4604 | 4469 |
| Prostate | Normal | 9614 | 0.06704 | 26 | 0.436 |  |  |
|  | Cancer | 8645 | 0.39908 | 108 | 0.697 | 4406 | 3629 |
| Skin | Normal | 7370 | 0.07258 | 97 | 0.578 |  |  |
|  | Cancer | 9841 | 0.51385 | 29 | 0.620 | 5164 | 4098 |
| Testis | Normal | 6825 | 0.06484 | 93 | 0.630 |  |  |
|  | Cancer | 10174 | 0.58501 | 73 | 0.747 | 5605 | 4128 |
| Thyroid | Normal | 14095 | 0.07634 | 50 | 0.477 |  |  |
|  | Cancer | 10259 | 0.27775 | 372 | 0.689 | 4509 | 4189 |
| Uterus | Normal | 11935 | 0.05935 | 91 | 0.542 |  |  |
|  | Cancer | 9956 | 0.95163 | 330 | 0.946 | 4778 | 3312 |

By examining only the fraction of intra-chromosomal links in cancer GCNs (Table 2), we identified two groups. The first group includes networks from bladder, breast, esophagus, kidney, liver, lung, ovary, and uterus, with more than 83% intra-chromosomal interactions. The second group consists of networks from brain, pancreas, prostate, skin, testis, and thyroid, with less than 66%. These two groups exhibit visually distinct structures when a force-directed layout is applied for network visualization (Supplementary Figure 4). In contrast, normal co-expression networks do not show such a division. The mean fraction of intra-chromosomal interactions in cancer co-expression networks is 0.7582 ± 0.2327, while networks from normal tissues have an average fraction of 0.0716 ± 0.0085, an order of magnitude lower.

**Fig. 4.**
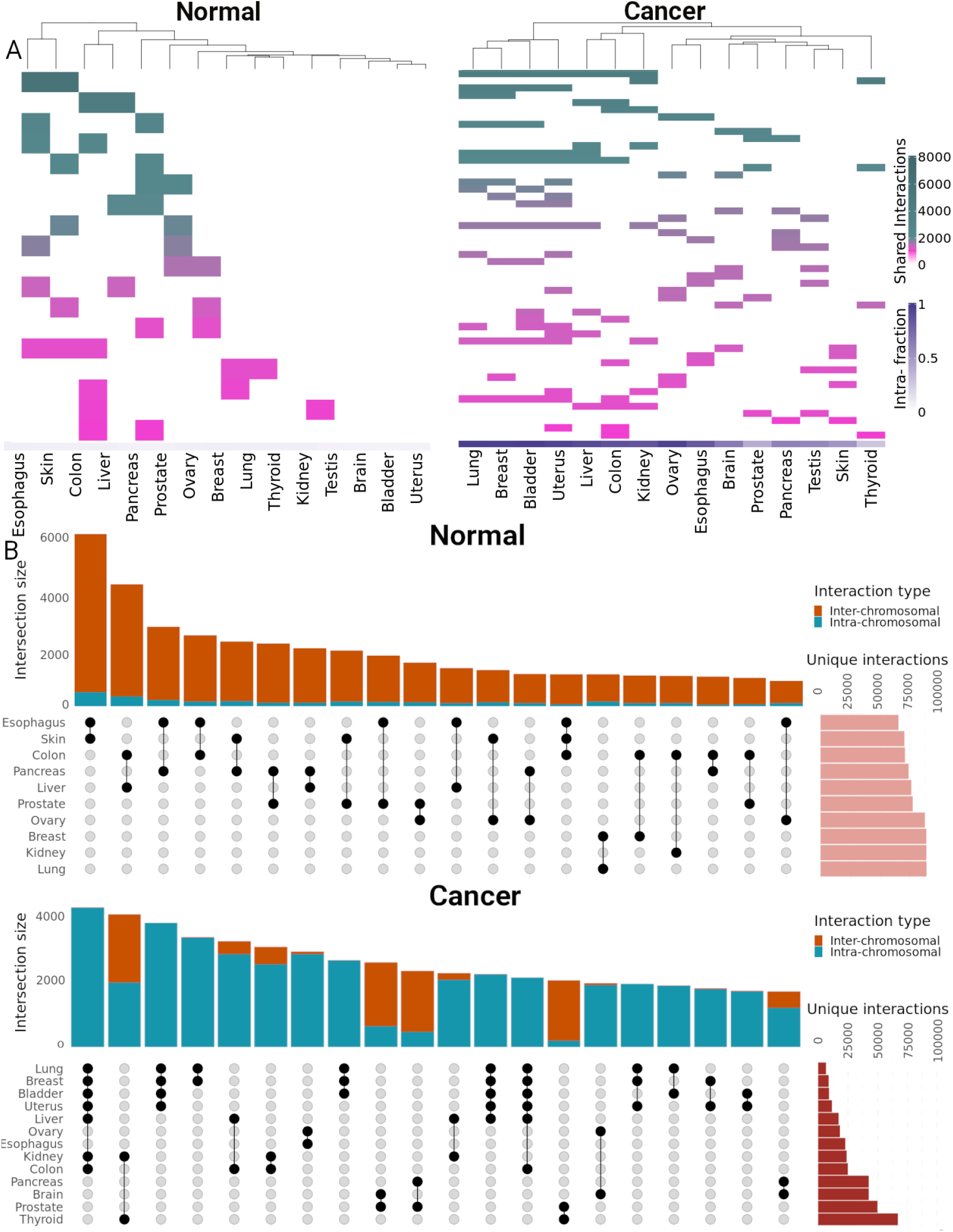
(A) Hierarchical clustering of interactions shared among tissues in normal and cancer co-expression networks, showing sets with more than 1,000 interactions. **(B)** Bar plots and upset blots displaying the top 20 sets of shared interactions. Normal co-expression networks exhibit a higher number of unique interactions, while cancer networks predominantly share intra-chromosomal interactions.

### Links are mostly unique in normal co-expression networks, but not in cancer

To compare co-expression networks at the interaction level, we performed hierarchical clustering based on the presence of gene-gene links across different tissues. Sets with more than 1000 shared interactions are shown in Figure 4A.

Clustering divides cancer GCNs into three major groups. Two clusters with a high fraction of intra-chromosomal links emerge: one includes breast, bladder, lung, and uterus, while the other consists of networks from colon, kidney, and liver cancer. The third group comprises networks with a lower fraction of intra-chromosomal links: ovary, esophagus, brain, prostate, pancreas, testis, skin, and thyroid. The largest set of shared interactions in the total set of GCNs occurs between normal networks in esophagus and skin.

Vertical bar plots in Figure 4B display the twenty sets with the highest number of shared links. Consistent with their composition, normal GCNs predominantly share interchromosomal interactions, while cancer networks share links on the same chromosome. Among cancer GCNs, lung, breast, and bladder have the fewest unique interactions, with 5,621, 7,894, and 8,607 respectively, whereas the thyroid cancer network has the most unique interactions, totaling 67,077.

Despite the visual similarity of normal networks in Figure 3 and their comparable fractions of intra-chromosomal links, they share fewer interactions among themselves than cancer networks, as shown in the heatmap (Figure 4A) and the horizontal bar plots in Figure 4B. The fifteen normal GCNs have an average of 83,642 unique interactions (±10, 362), while cancer GCNs have an average of 29,366 (±20, 379) unique interactions.

An interesting aspect related to the data source is highlighted in the vertical bar plots: the top 14 sets of shared interactions in the normal phenotype are exclusively from tissues in the UCSC Xena dataset. This pattern does not occur in cancer, where the largest set includes tissues from both data sources.

### Riboprotein genes constitute a cluster shared by cancer and normal co-expression networks

To identify coexpression patterns present in both normal and cancer GCNs across multiple tissues, we constructed a common network featuring the most frequent interactions, defined as those occurring in more than 10 tissues. These networks are displayed in Figure 5. The cancer network contains 12,802 links, whereas the normal network has only 355 edges. The intersection of normal and cancer networks includes 92% of the genes from the common normal network and 59% of its links. Most clusters in the cancer common network consist of genes from the same chromosome, although small subsets of nodes are formed by inter-chromosomal interactions.

**Fig. 5.**
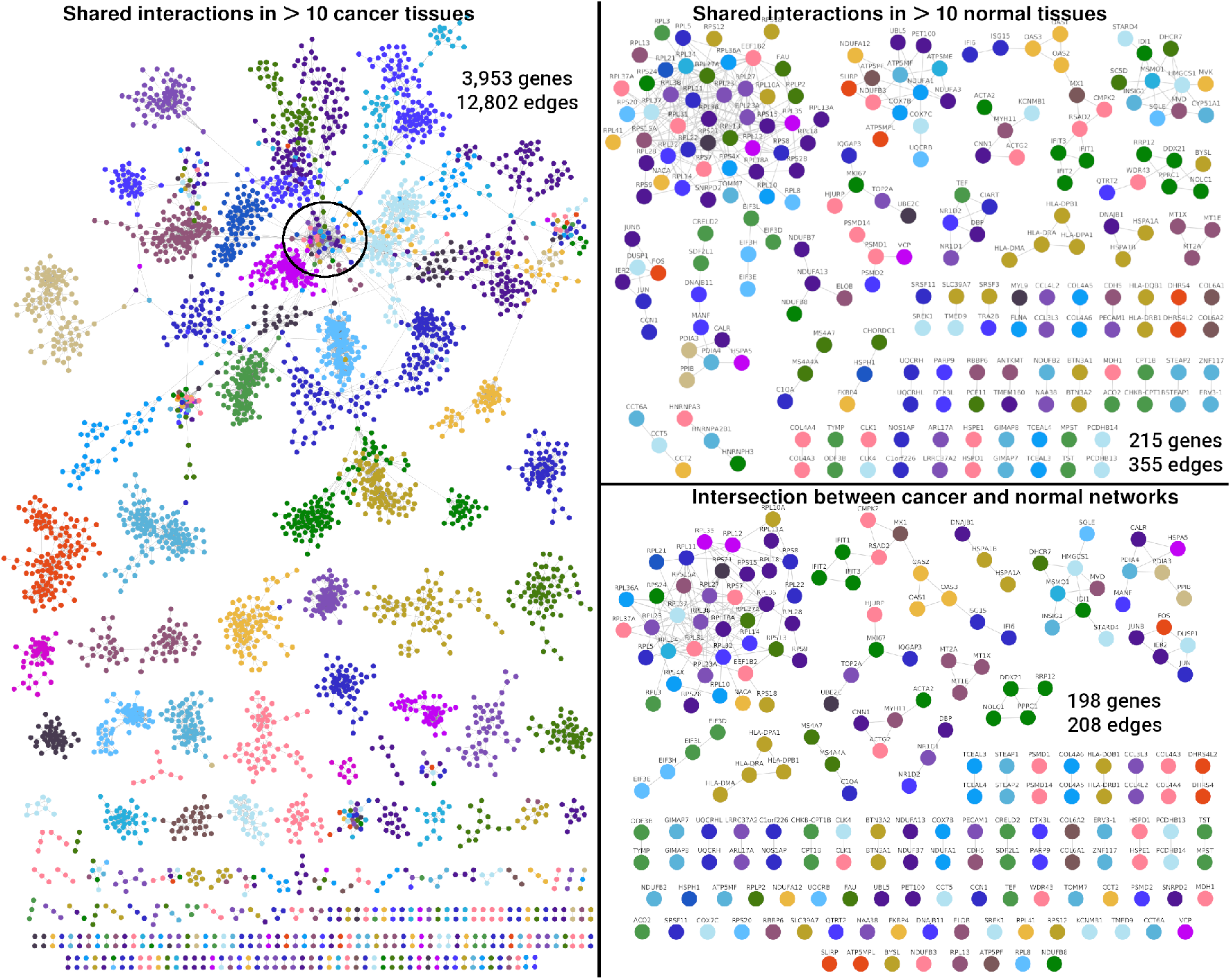
Cancer and normal common networks, constructed from interactions shared across more than ten tissues, along with their intersection, visualized using a forcedirected layout. Genes encoding ribosomal proteins, connected by inter-chromosomal links,form the largest component in the intersection. This component is highlighted with a circle at the center of the largest component in the cancer network.

The largest connected component in the intersection of normal and cancer networks comprises thirty-nine genes, located at the center of the giant component of the cancer network (circled in black in Figure 5). Thirty-seven of these genes encode molecules in the ribosomal protein family, including members of both the small and large subunits. The remaining two genes are EEF1B2, which is involved in transferring aminoacylated tRNAs to the ribosome (33), and NACA, which prevents the mistranslocation of aberrant nascent proteins to the endoplasmic reticulum (34). The presence of this cluster indicates that ribosomal genes maintain a pattern of high co-expression among themselves in all analyzed tissues and under both conditions, cancerous and normal.

### The community structure of co-expression networks is associated with biological processes

The forcedirected layout applied to the co-expression networks for visualization (Figure 3 and Supplementary Figure 4) suggests a modular structure influenced by chromosomal gene location, particularly in cancer GCNs with a higher fraction of intrachromosomal links. To explore this further, we performed a community structure analysis using the Louvain algorithm (35). This algorithm has been shown to effectively identify biologically relevant network structures (36, 37). Table 2 lists the number of communities identified in each network.

For each community, we calculated its chromosomal assortativity, a nominal assortativity to quantify the community’s bias towards a single chromosome. The distributions of chromosomal assortativities for cancer and normal co-expression networks are presented as violin plots in Figure 6. These plots show that network communities in cancer tend to have values close to 1, indicating that genes in a community mostly belong to a single chromosome. In contrast, normal GCNs exhibit an opposite trend, with chromosomal assortativity values clustering towards -1 in tissues such as bladder, breast, liver, ovary, prostate, thyroid, and uterus, and values distributed between -1 and 1 in other tissues. This distribution suggests either a link bias towards different chromosomes or mixed trends, respectively.

**Fig. 6.**
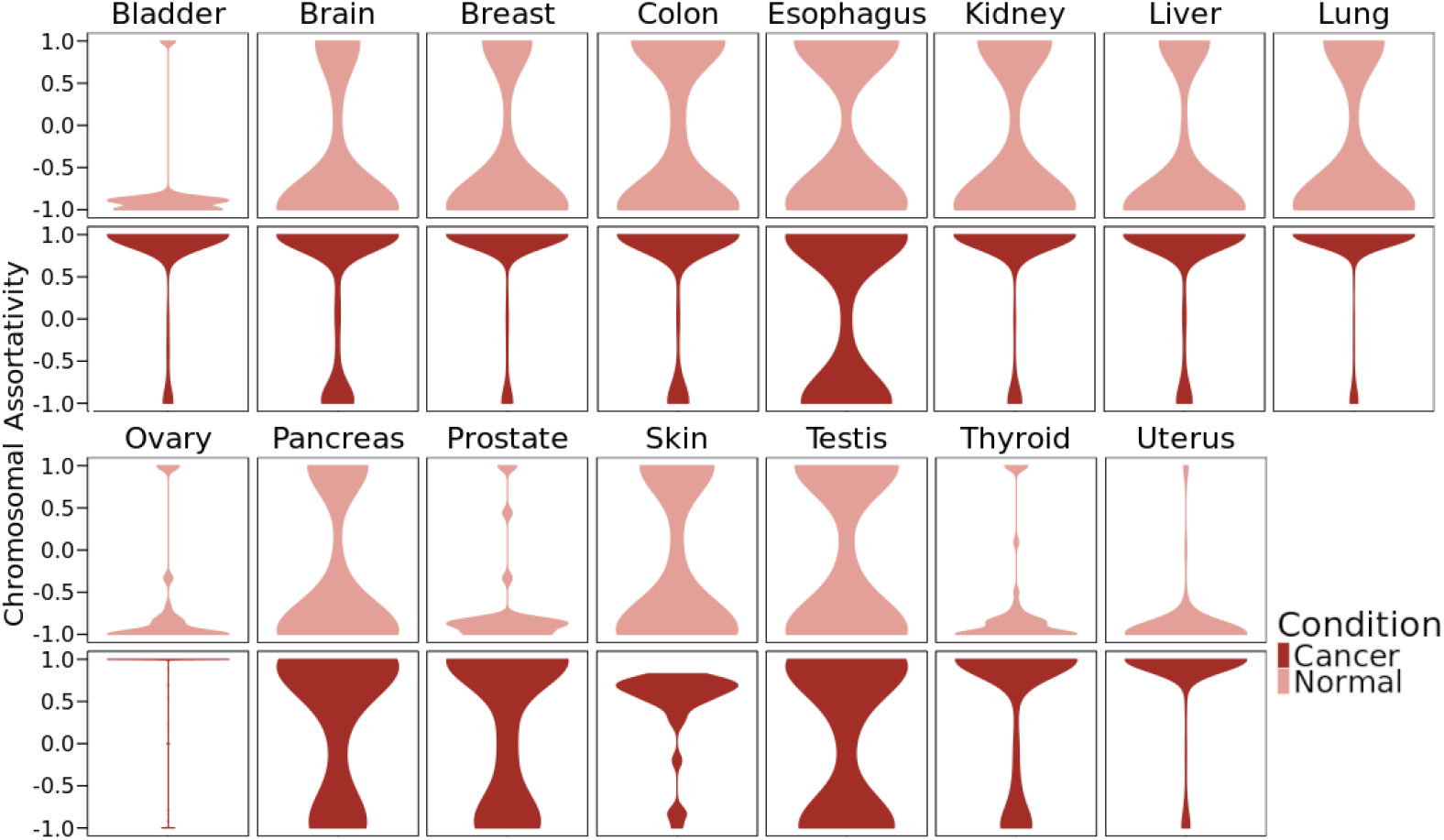
Chromosomal assortativity in network communities of all fifteen tissues analyzed. Normal tissues tend towards values near -1, indicating communities are primarily composed of inter-chromosomal links. In contrast, cancer tissues show assortativity values closer to 1, reflecting communities dominated by intra-chromosomal interactions.

#### Shared GO processes in co-expression networks’ communities

We conducted Gene Ontology (GO) over-representation analyses to identify biological processes associated with each community in GCNs. By linking communities from the fifteen networks to their enriched biological processes, we created a bipartite network for both normal and cancer phenotypes. We then selected processes with a high degree (D > 10) to identify the most common GO processes in each phenotype. The resulting networks are displayed in Figure 7 and Figure 8.

**Fig. 7.**
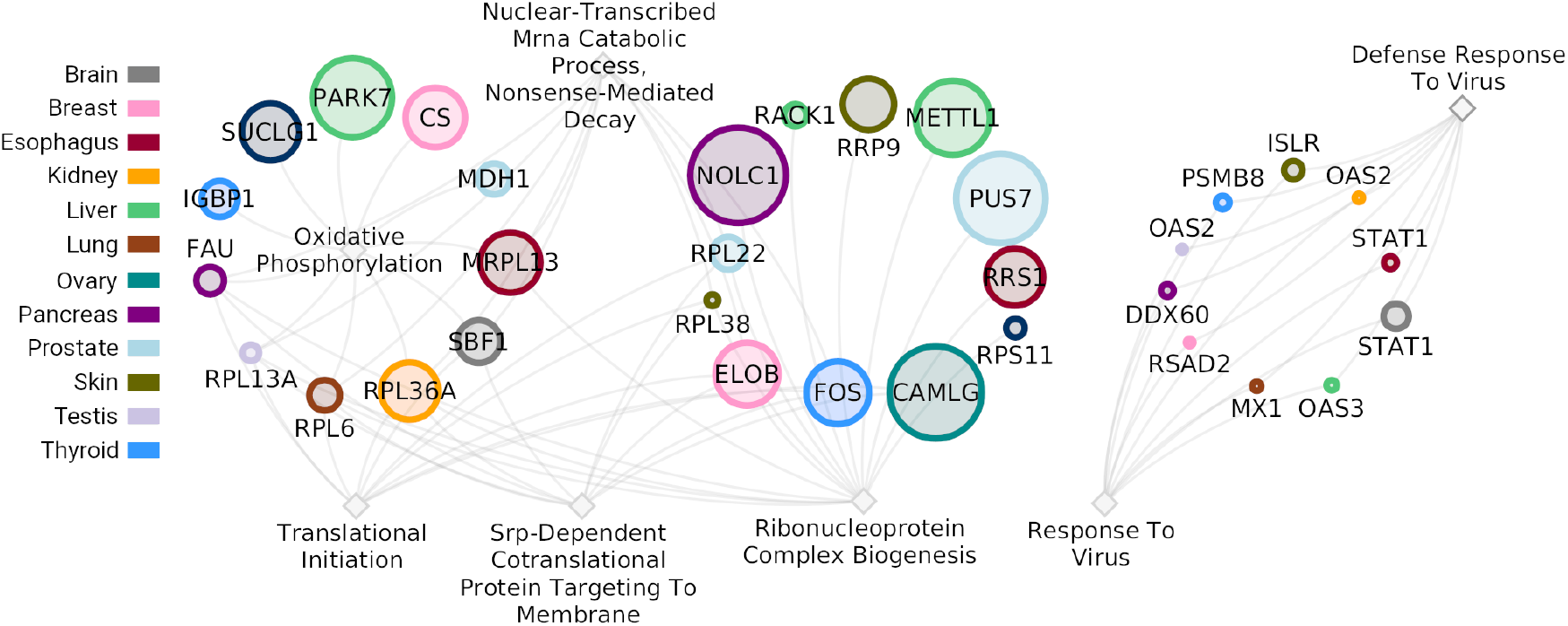
Bipartite network of the most common Gene Ontology biological processes with a degree greater than ten, joined to their communities in the normal phenotype. Diamonds represent enriched processes and circles represent communities colored according to their tissue.

**Fig. 8.**
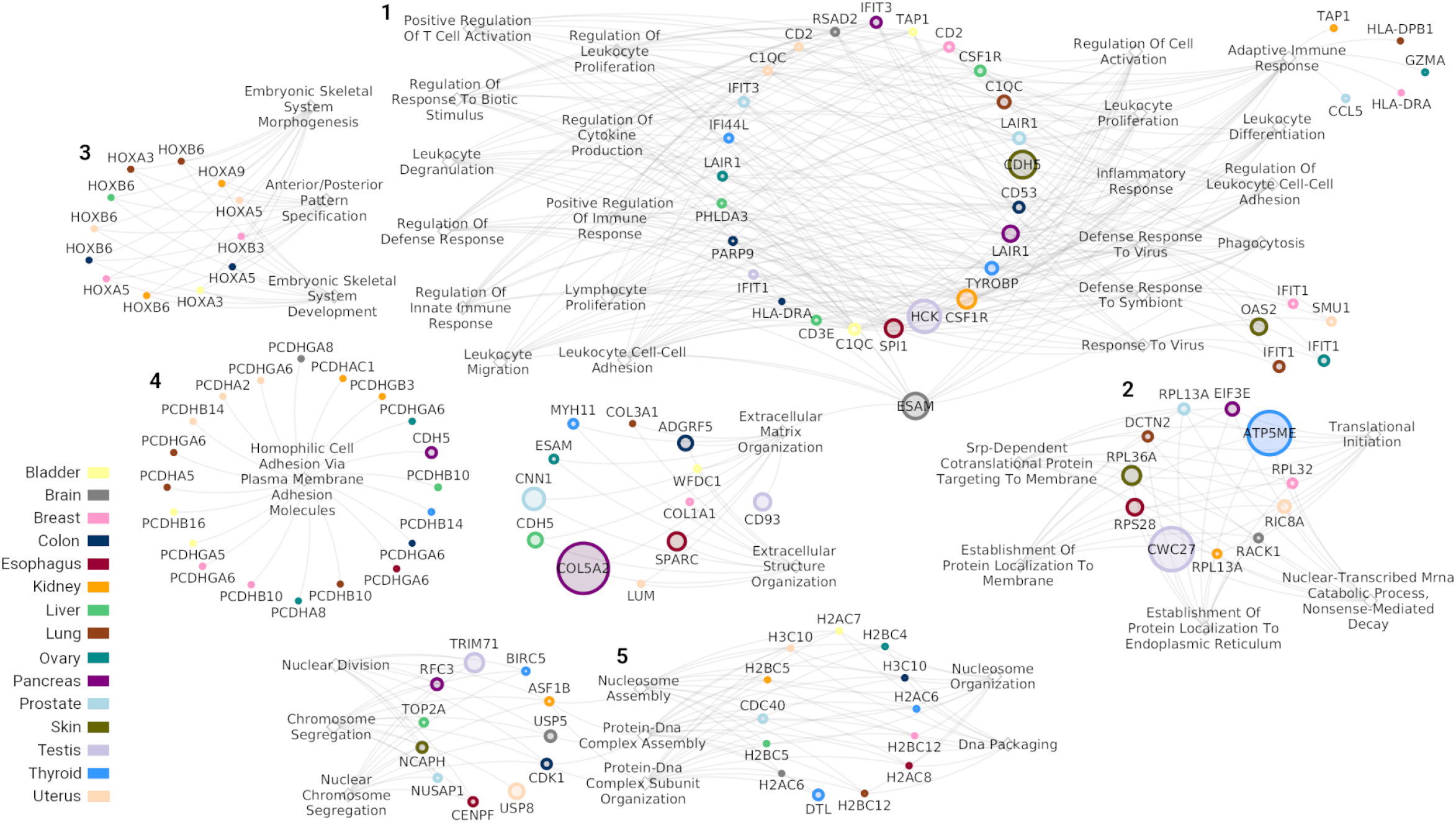
Bipartite network of the most common Gene Ontology biological processes, with a degree greater than ten, joined to their communities in the cancer phenotype. Diamonds represent enriched processes and circles represent communities colored according to their tissue.

In normal networks (Figure 7), common GO processes generate two connected components. The largest includes terms related to translation and protein localization, such as *ribonucleoprotein complex biogenesis* and *translational initiation*, while the other component contains terms related to response to virus. We found no common GO processes in bladder and uterus enriched GO terms.

In cancer networks (Figure 8), the bipartite subnetwork of common GO processes forms five connected components. The largest connected component (identified with the number 1) is related to immune response and includes the process most commonly found in cancer networks: *adaptive immune response*. It is enriched in twenty-one communities from all cancer types, except for skin cancer. Some of these communities include genes in chromosome 6, encoding almost all members from the major histocompatibility complex II, such as HLA-DP, HLA-DM, HLA-DOA, HLA-DQ, and HLA-DR. Other processes include *positive regulation of T cell activation, lymphocyte proliferation*, and *inflammatory response*.

We also find one component that mirrors the ribosome-associated processes in normal networks (number 2), with terms like *translational initiation* and *establishment of protein localization to the endoplasmic reticulum*, indicating high co-expression among ribosomal genes. A third module (3) includes twelve communities with HOXA and HOXB gene families, linked to processes like *embryonic skeletal system morphogenesis*. These communities are entirely intrachromosomal, located on chromosomes 7 (HOXA) and 17 (HOXB).

The PCDHG and PCDHB protocadherin clusters form another component (4), associated with *homophilic cell adhesion via plasma membrane adhesion molecules*. This cluster includes communities from twelve cancer types, such as bladder, breast, kidney, ovary, and thyroid. Most communities are intra-chromosomal, with genes on chromosome 5 showing underexpression, particularly in bladder, breast, and uterus, as previously reported in Luminal A breast cancer (29).

The last component (5) is associated with histone-related activities, featuring GO processes such as *nucleosome assembly* and *chromatin organization involved in negative regulation of transcription*, involving genes from the H2A, H2B, H3C, H4C, H1, and H4 families.

#### Normal- and cancer-only associated processes

The bipartite networks help identify processes unique to either normal or cancer co-expression networks. Sets of similar normal-only processes are shown in Figure 9, and cancer-only ones in Figure 10. Again, diamonds represent enriched processes in network communities (*p*_*adj*_ *<* 1*e*^*−*10^), while colored circles represent communities in their respective tissues.

**Fig. 9.**
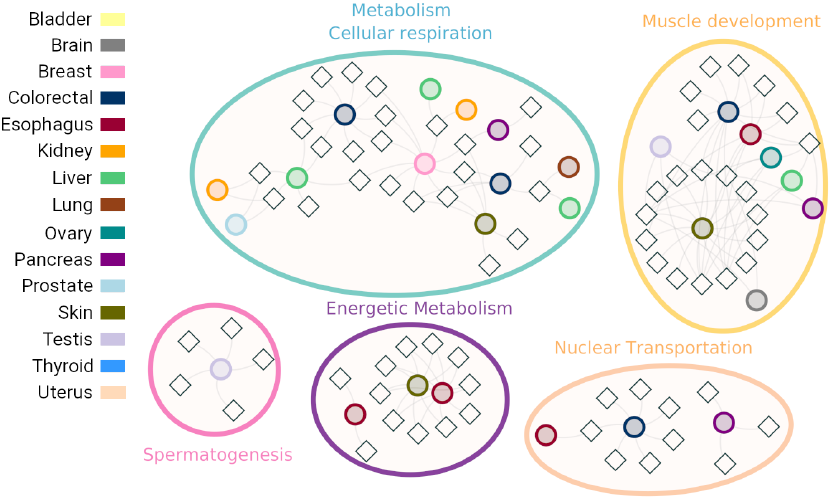
Bipartite network of Gene Ontology processes and their associated communities found exclusively in the normal phenotype.

**Fig. 10.**
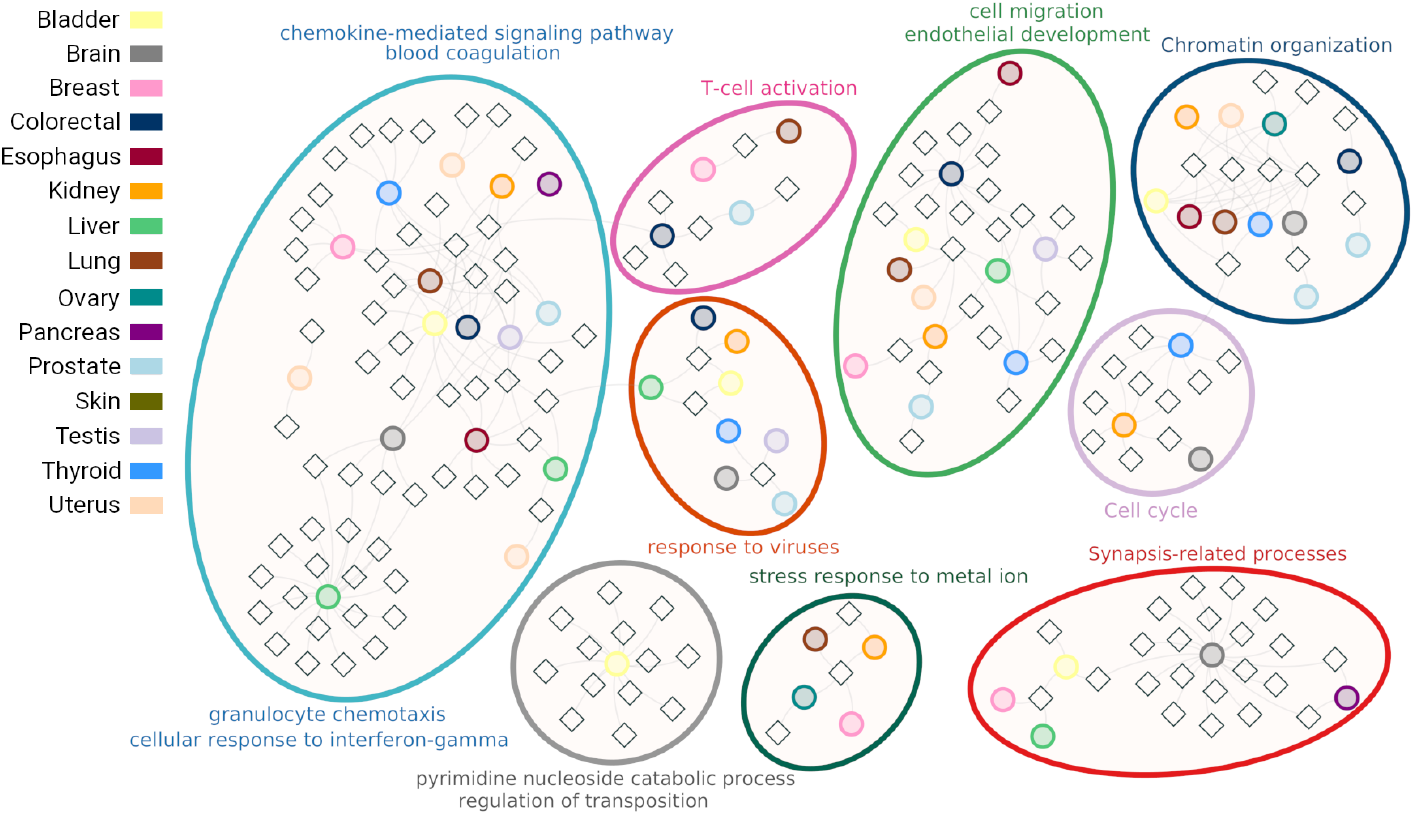
Bipartite network of Gene Ontology processes and their associated communities found exclusively in the cancer phenotype. Similar GO terms were grouped. Diamonds represent enriched processes and circles represent communities colored according to their tissue.

In normal network communities, we identified five sets. The largest, related to metabolism and cellular respiration, includes communities from eight tissues with processes like *mitochondrial translation* and *regulation of lipid metabolic*. The muscle development set also spans eight tissues, featuring processes such as *cardiac cell development* and *myofibril assembly*. The remaining sets are *spermatogenesis, energetic metabolism*, and *nuclear transportation*, all essential for cell maintenance.

The largest set of cancer-only processes is linked to immune response, involving communities from thirteen tissues and processes like *antigen processing and presentation, interleukin-1 production, lymphocyte differentiation*, and *response to chemokine*, among others. Two additional sets relate to immune response activation: *T-cell activation* and *response to viruses*. The network also includes processes altered in cancer, such as *cell migration, cell cycle*, and *chromatin organization*.

In these networks, we identified GO terms uniquely associated with specific tissues, highlighting individual features of their co-expression networks and linking them to known tissue-specific biological processes. For instance, the *spermatogenesis* set in the normal network includes a single community from testis, where the gene with the highest pagerank is Boll or BOULE, essential for sperm development (38). This community is associated with processes like *cilium movement, axoneme assembly, spermatogenesis*, and *single fertilization*. In the skin normal co-expression network, enriched terms in the MLANA community include *melanin biosynthetic process, melanin metabolic process*, and *pigment biosynthetic process*. The MLANA or Melan-A gene, encodes a protein involved in melanosome biogenesis (39).

Conversely, in the brain GCN, the CCKBR community is related to processes such as *neurotransmitter secretion, synaptic signaling*, and *regulation of membrane potential*, among others. The 257 genes involved in the CCKBR community enrichments exhibit an underexpression trend, with an average *log*_2_ fold change of *−*0.703.

### The loss of inter-chromosomal co-expression is not present in other diseases

To determine if the loss of inter-chromosomal co-expression is a phenomenon exclusively present in cancer, we analyzed two RNA-Seq datasets downloaded from the Gene Expression Omnibus database. GSE174367, a study of late-stage Alzheimer’s Disease (AD) (40), with 44 AD samples and 46 control samples from brain tissue and GSE184050, a Type-2 Diabetes (T2D) study (41) with 50 blood T2D samples and 66 blood control samples. We calculated mutual information between gene pairs using the same algorithm as for cancer datasets and extracted the top 100,000 interactions to construct co-expression networks. In these diseases, the fraction of intra-chromosomal interactions in co-expression networks is similar for both normal and disease phenotypes, as sown in Figure 11. For AD, the intra-chromosomal fractions are 0.060 for normal and 0.0558 for disease networks. In the T2D network, these fractions are 0.0733 and 0.0734, for normal and disease respectively.

**Fig. 11.**
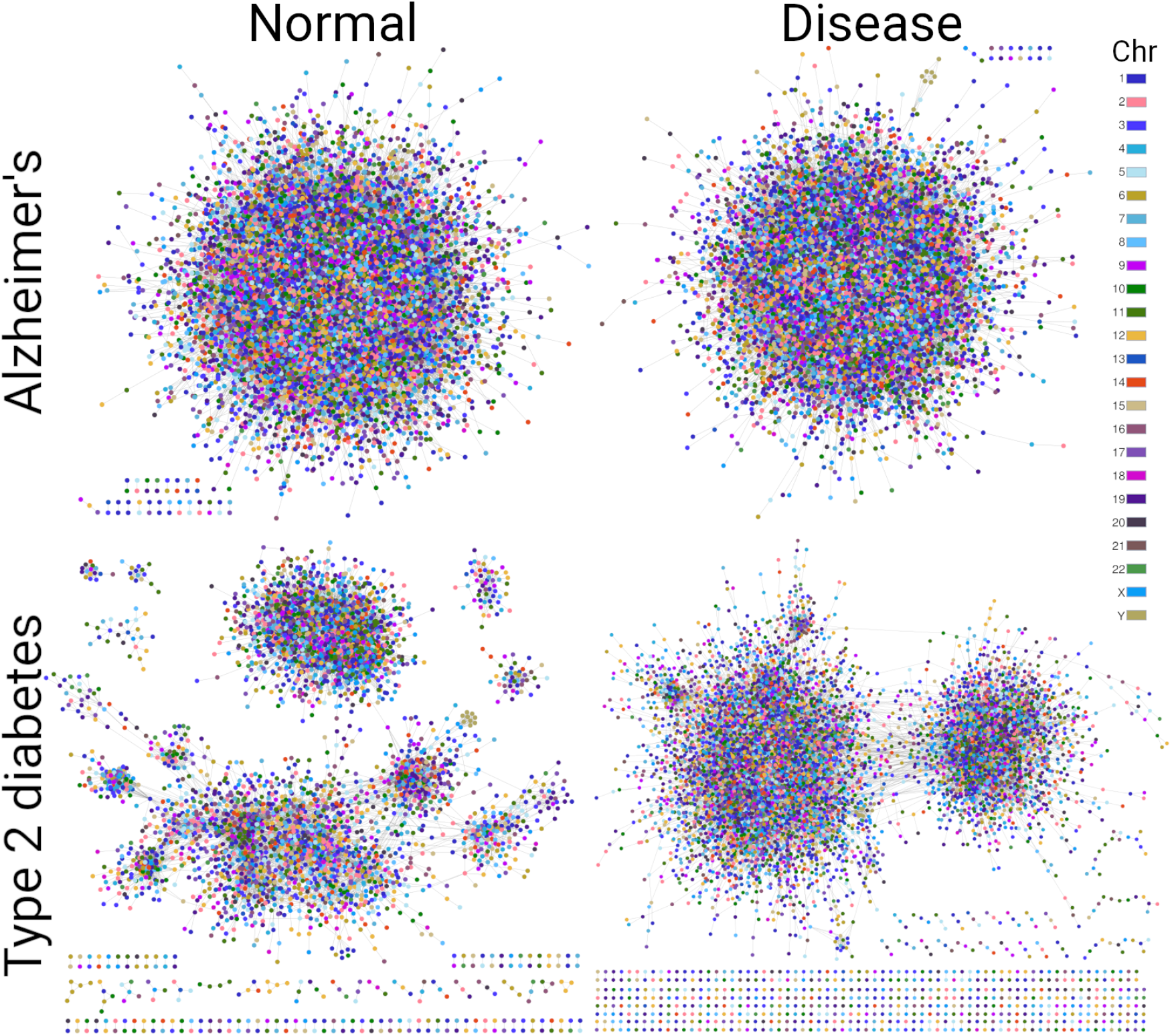
Co-expression networks from Alzheimer’s disease and type-2 diabetes composed by the top 100,000 MI interactions.

None of these networks present chromosome associated gene interaction patterns, and for both phenotypes, normal and disease networks have similar structures. These findings indicate that the loss of inter-chromosomal co-expression is not observed in these chronic degenerative diseases.

## Discussion

In this study, we characterized two cancer-associated phenomena: the loss of inter-chromosomal co-expression and the distance-dependent decay of gene co-expression for intrachromosomal interactions. co-expression profiles, calculated using Mutual Information (MI), revealed that the strongest correlation values occur between genes located on the same chromosome. Furthermore, by linking intra-chromosomal interactions to the physical distance between genes (measured in base pairs), we found that, in cancer, high correlation values are concentrated in close proximity. In contrast, normal tissues (i.e., healthy adjacent tissue used as control in TCGA and samples tagged as Normal Tissue in GTEx-Xena) do not exhibit a bias towards intra- or inter-chromosomal interactions or any specific physical distance between gene pairs, except for very close distances, as illustrated in the left side of the distance plots for normal tissue profiles (Figure 2 and Supplementary Figure 2).

The difference between cancer and normal profiles regarding the strength of gene co-expression and intra-chromosomal distance is evident in Figure 2 and Supplementary Figure 2. All cancer tissues exhibit a significant decay in the mean MI value relative to the base pair distance between same-chromosome genes. Conversely, MI values in normal networks reach a plateau early in the plots. In both cases, standard deviation profiles support these findings.

Previous studies have shown high local gene co-expression in normal tissues for genes located near each other (within 1 Mb), regardless of strand or transcription orientation (24, 25, 27, 28), indicating that gene transcription is influenced by physical location. Evidence suggests that transcription contributes to gene cluster formation and fine-scale chromatin organization (42, 43). In cancer profiles, highly co-expressed gene pairs span greater distances, with some reaching a decay plateau around 50 Mb.

The assessment of the contribution of different regulatory elements such as enhancers, transcription factor binding sites, CTCF binding sites, and Hi-C contacts, to the local gene co-expression in normal tissues done by (23) indicated that no feature can uniquely explain the co-expression patterns observed; rather, they contribute complementarily. Our previous work showed that intra-chromosomal communities in Luminal A and B breast cancer are not bound by CTCF sites, and copy number alterations do not affect coexpressed communities with similar differential expression trends (29, 44, 45). We also explored epigenetic mechanisms, particularly miRNA regulation, which alters co-expression in genes related to epithelial-mesenchymal transition (46, 47), promotes subtype-specific pathways in breast cancer (48), and is linked to progression in clear cell renal carcinoma (49). However, these examples involve inter-chromosomal interactions, highlighting the need for further research on regulatory mechanisms enabling local co-expression in normal tissues and alterations expanding co-expression regions in cancer.

It is worth recalling that our MI calculations compare gene expression vectors independently, without influence from chromosomal position or phenotype. Thus, the higher statistical dependency among neighboring intra-chromosomal genes in cancer suggests a phenomenon enhancing close expression patterns across long chromosomal regions. We hypothesize that decay behavior in cancer could be related to functional or mechanical features of the transcriptional machinery impacting gene transcription.

Although all fifteen cancer profiles show a loss of long-range co-expression, Figure 2 and Supplementary Figure 2 reveal differences in extent among tissues. For example, thyroid and prostate profiles show less pronounced differences, while bladder, breast, lung, and ovary exhibit greater distances between intra-chromosomal interactions and more pronounced decay, possibly reflecting tissue-specific cancer manifestations. Thyroid GCN shows the highest number of unique interactions and it lacks some common features from other networks, such as the presence of a community of HOX genes. According to GLOBOCAN, thyroid cancer has one of the lowest mortality rates (50) and the most common subtypes, papillary and follicular, have a 5-years survival rate of almost 100% (51, 52). We have shown that in papillary and follicular thyroid cancer, relevant pathways are significantly less deregulated than in the most aggressive subtypes (53). Therefore, the particular features of the thyroid co-expression profile may reflect its less aggressive nature. Further research into subtype-specific networks could be valuable.

Normal and cancer GCNs differ visually, especially those with higher intra-chromosomal link fractions: bladder, breast, esophagus, kidney, liver, lung, ovary, and uterus (Table 2 and Supplementary Figure 4). Previous studies have reported co-expression clusters and functional associations (54–56), as well as connectivity loss in cancer networks derived from microarray data (57). However, the loss of longrange interactions and gain of close-distance links in had not been described as prevalent properties of cancer GCNs.

When comparing intersections among networks, a clear difference emerges between cancer and normal phenotypes: normal GCNs share fewer interactions than cancer networks. We previously reported this for lung cancer, where squamous cell lung carcinoma and lung adenocarcinoma networks shared more interactions than their normal counterparts (20). The co-expression program is a manifestation of the cell state, and the similarity among cancer tissues from various origins suggests functional parallels. We propose that the extensive set of shared intra-chromosomal interactions is linked to cell dedifferentiation processes during tumor development (58–60), closely related to phenotypic plasticity, one of the newly described hallmark capabilities (6).

Despite the small number of connections shared among normal co-expression networks, a significant cluster of riboprotein-encoding genes forms the largest component in the common normal tissue network (Figure 5). This module also appears in the common cancer network, indicating that ribosomes maintain a high co-expression pattern in both normal and cancer tissues. As essential cellular components, ribosomal genes are highly conserved across species (61), which may explain their prevalence in co-expression networks. Their presence underscores the importance of the translation process and it exhibits the capability of coexpression networks to capture important characteristics associated with a phenotype.

Regarding network structure, cancer GCNs exhibit observable modular behavior (Supplementary Figure 4), with higher modularity values than normal networks (Table 2), particularly in networks with high intra-chromosomal fractions. This suggests that, in normal phenotypes, genes interact in a less compartmentalized manner, with transcriptional events driven by cellular functions and environmental signals. In contrast, segregated modules in cancer networks may indicate a partial loss of information spread or a gain in specific co-expression regions.

Chromosomal assortativity (Figure 6) serves as a metric to summarize a key phenomenon: normal networks communities tend towards inter-chromosomal interactions, while cancer communities predominantly consist of intrachromosomal links. Furthermore, no networks exhibit chromosomal assortativity values near zero, indicating communities are formed from either intra- or inter-chromosomal links. The modular composition, distance-associated co-expression decay, and high chromosomal assortativity in network communities suggest highly orchestrated co-expression events involving nearby genes in cancer GCNs. The loss of longdistance communication at the transcriptomic level and the emergence of co-expression hotspots may contribute to the disruption of essential cellular functions.

This is further illustrated in bipartite networks of Gene Ontology processes shared across more than ten tissues, whether normal or cancer, and in the networks of normal-only and cancer-only processes (Figure 7, Figure 8, Figure 9, Figure 10). GO terms in normal networks are associated with cellular functions such as translation, metabolism, and development. In contrast, while inter-chromosomal cancer communities show a notable presence of immune response-associated processes, most GO processes are enriched in same-chromosome communities comprising gene families such as HOXA and HOXB, protocadherins, histone variants, and metallothioneins.

These gene families are known to be altered in different cancer types. HOX genes are implicated in the formation of cancer stem cells (62–64) contributing to both hematologic malignancies and solid tumors. They are also associated with the dedifferentiation process and phenotypic plasticity (6). Protocadherins may exhibit low expression due to long-range epigenetic silencing via hypermethylation, which promotes migration and proliferation in various cancers (65–67). Metallothioneins display diverse expression patterns that affect tumor growth, microenvironment remodeling, and drug resistance (68, 69). Histones play a crucial role in the regulation of chromatin structure and are involved in cancer development and progression (70), with aberrant expression of certain histone variants reported in multiple cancer types (71).

In addition to modules shared among multiple networks, some GCNs reveal tissue-specific associations. For example, the brain cancer co-expression network presents a set of neural system-related processes with predominantly underexpressed genes. This suggests an association among differential gene expression, gene co-expression, and functional features of a specific network community, demonstrating the ability of GCNs to capture phenotype-specific manifestations.

The affirmation that loss of long-range co-expression is specific to cancer is supported by results presented in Figure 11. Two chronic-degenerative diseases such as Alzheimer’s disease and Type-2 Diabetes present strong similarities in their co-expression landscapes, both visually and in chromosomal assortativity values. Evaluating the presence of this phenomenon in other diseases requires additional disease-specific datasets.

Despite their unique features, our results are consistent across fifteen cancer tissues compared to their normal counterparts. There are no significant differences related to data source; both TCGA and UCSC Xena datasets include tissues with varying levels of distant co-expression loss. Moreover, the number of samples does not appear to influence network structure, as previously assessed in breast (16) and lung cancer (20). However, there are some considerations that need to be addressed in future analyses. For instance, the hierarchical clustering shown in Figure 4 does not group according to tissue similarity or cancer origin. Further evaluation of the nature of the displayed similarities is needed. In that same figure, the biggest intersection sets which include esophagus, skin, and pancreas interactions, belong to the UCSC Xena dataset, with data from GTEx; we should evaluate whether the pre-processing steps added some bias.

Our findings offer new insights into the organization of the cancer transcriptome. Future research could build on these results by incorporating additional datasets, such as singlecell RNA-Seq, to help identify the specific cell types driving this phenomenon. Integrating multi-omics datasets would further clarify the role of altered regulatory mechanisms in the disruption of distant gene co-expression. Targeted experimental studies are also crucial to unravel the molecular processes underlying this alteration. The loss of long-range co-expression in cancer phenotypes may reflect a fundamental reorganization of transcriptional regulation, representing a common and distinctive feature that warrants deeper exploration.

## Methods and Materials

### Databases

RNA-Seq data from The Cancer Genome Atlas (30) was downloaded from the Genomic Data Commons (GDC) Data Portal for bladder, breast, kidney, lung, thyroid and uterus tissues in *STAR-Counts* format. Cancer datasets included samples tagged as Primary Tumor, while normal datasets included samples tagged as Solid Tissue Normal. Gene expression *RSEM expected counts* were downloaded from the UCSC Xena browser (31), which integrates TCGA and GTEx (72) data using the Toil RNA-seq pipeline (32). For brain, colon, esophagus, liver, ovary, pancreas, prostate, skin, and testis, normal counts were obtained from GTEx-Xena, tagged as Normal Tissue, while cancer counts were identified as Primary Tumor. RSEM expected counts were transformed from log2+1 to integer count values. Sample counts per tissue and condition are shown in Table 1.

### Data processing

The RNA-Seq data processing and differential expression analysis pipeline was adapted from (16). A Snakemake (73) workflow was implemented to create a reproducible analysis: https://github.com/ddiannae/llrc-pipeline. A second pipeline was created for the analysis of co-expression profiles and co-expression networks: https://github.com/ddiannae/llrc-networks.

### Pre-processing and normalization

Pre-processing was performed per tissue and sample type. TCGA samples were annotated using GENCODE v36 (https://gdc.cancer.gov/about-data/gdc-data-processing/gdc-reference-files). Only protein-coding genes, conventional chromosomes, and genes present in GENCODE v44 (July 2023) were retained. After assembling raw count matrices, datasets from both sources followed the same pipeline.

The `NOISeq` R library (74) was used for quality control. Genes with mean expression *<* 10 or zero counts in more than 50% of samples were removed. Gene counts expression boxplots revealed that some samples from the XENA database had extremely low values; hence, samples with mean expression values two standard deviations above or below the total mean were excluded.

Gene counts were transformed to transcripts per million (TPM). Principal Component Analysis plots revealed overlap between normal and tumor samples, thus `ARSyNSeq` (75) was applied for multidimensional noise reduction using default parameters.

### Mutual Information (MI) calculation

The ARACNE algorithm (76) was used to calculate mutual information (MI), quantifying statistical dependence between gene pairs. MI values were computed per tissue and sample type. A `Singularity` container was created to run ARACNE within the Snakemake workflow (https://github.com/ddiannae/ARACNE-multicore).

### Differential expression analysis

Differentially expressed genes in cancer samples were identified using the `DESeq2` (77) R package. P-values were adjusted for multiple comparisons using the False Discovery Rate (FDR), with *p*_*adj*_ *<* 0.05 as the threshold and no threshold was applied to log fold change (logFC) values.

### Co-expression profile analysis

#### Identification of intra-chromosomal fraction

Gene pairs were ranked by MI in descending order, and the fraction of intra-chromosomal interactions was calculated at thresholds ranging from 1,000 to 100 million top MI pairs. Kolmogorov-Smirnov tests were used to compare intrachromosomal fractions between cancer and normal phenotypes.

#### Distance versus MI analysis

Intra-chromosomal gene pairs were grouped into bins of 1,000 interactions based on base pair distance. For each bin, mean MI, mean distance, and MI standard deviation were calculated. Wilcoxon rank sum tests were used to compare MI values between bins.

### Network Analysis

#### Networks and community detection

co-expression networks were constructed using the top 100,000 MI interactions per tissue and sample type. Communities were detected using the Louvain algorithm (35) implemented in `igraph` (78), with MI values as link weights.

#### Nominal assortativity

Chromosomal assortativity was calculated by dividing the difference between intraand interchromosomal links by the total number of links in a community.

#### GO Overrepresentation Analysis

We used the `enrichGO` function from the `clusterProfiler` R package (79) to identify over-represented Biological Process terms in Gene Ontology (GO) (80). Enrichment analysis was performed for communities with at least five genes, retaining GO terms with a minimum size of ten. The universe set was defined by the genes in the original expression matrix for each tissue. GO terms with *p*_*adj*_ *<* 1*E*^*−*10^, adjusted using the BenjaminiHochberg method, were considered significant.

## Supplementary Figures

1. Fraction of intra-chromosomal interactions at different thresholds for the fifteen tissues in the study.
2. Average MI values of bins of one thousand intrachromosomal interactions plotted against base pair distance for all co-expression profiles.
3. P-values of Wilcoxon rank sum tests comparing the distribution of MI values for bins of intra-chromosomal interactions for all co-expression profiles.
4. Co-expression networks from the top 100 thousand MI interactions for normal and cancer tissues.

## Supplementary Files

1. Kolmogorov-Smirnov p-values for the comparison between fraction of intra-chromosomal interactions in normal and cancer tissues at different MI thresholds.

## Author Contributions

DGC performed computational analyses, developed and implemented programming code, performed pre-processing and low-level data analysis, made the figures, drafted the manuscript. EHL developed the theoretical approach, supervised the statistical analysis and reviewed the manuscript. JEE conceived and designed the project, supervised the project, made the figures, drafted and reviewed the manuscript. All authors read and approved the final version of the manuscript.

