## Supplementary figures and images for "Loss of long-range co-expression is a common feature in cancer"

### Supplementary Figure 1

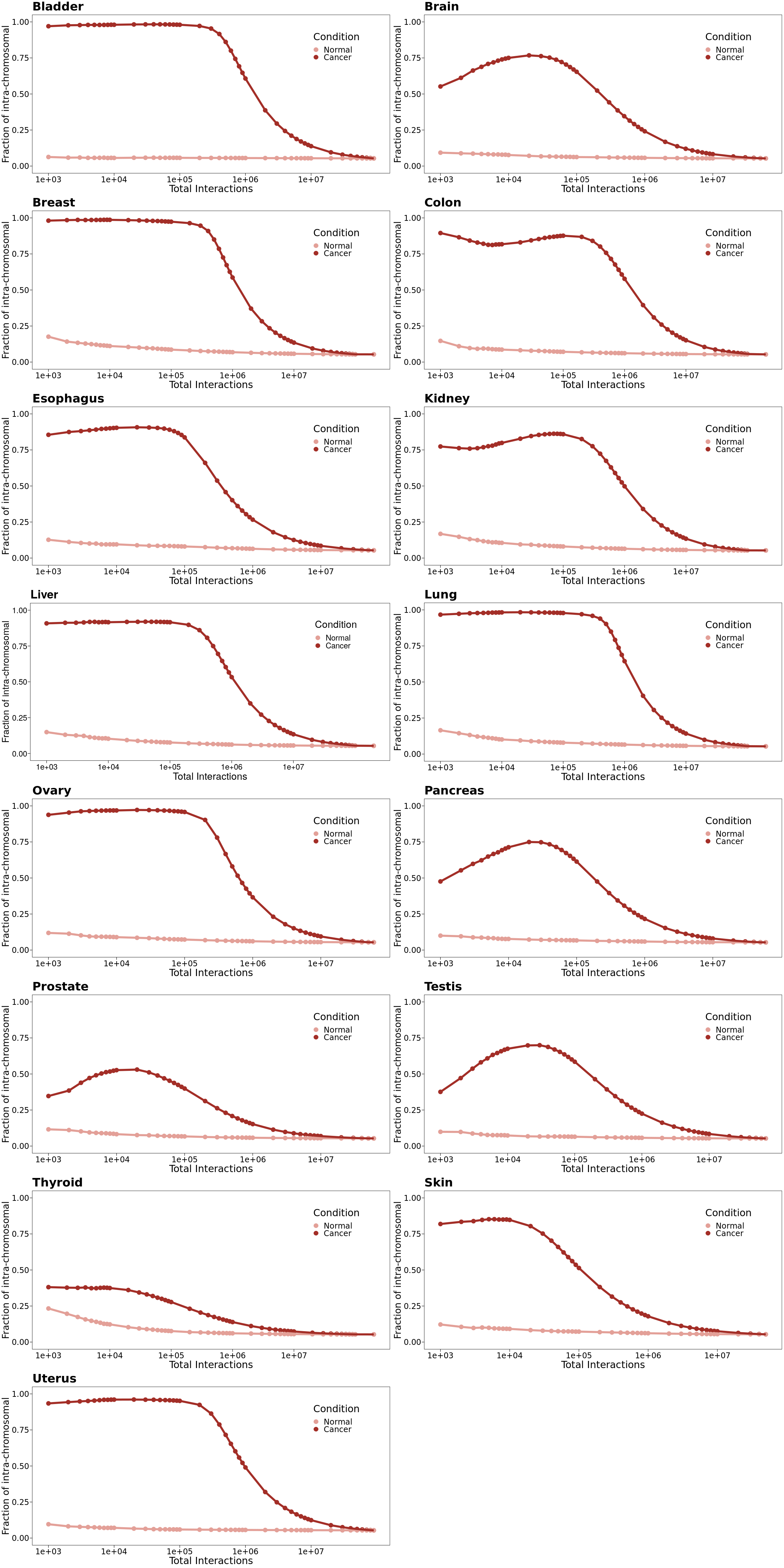

### Supplementary Figure 2

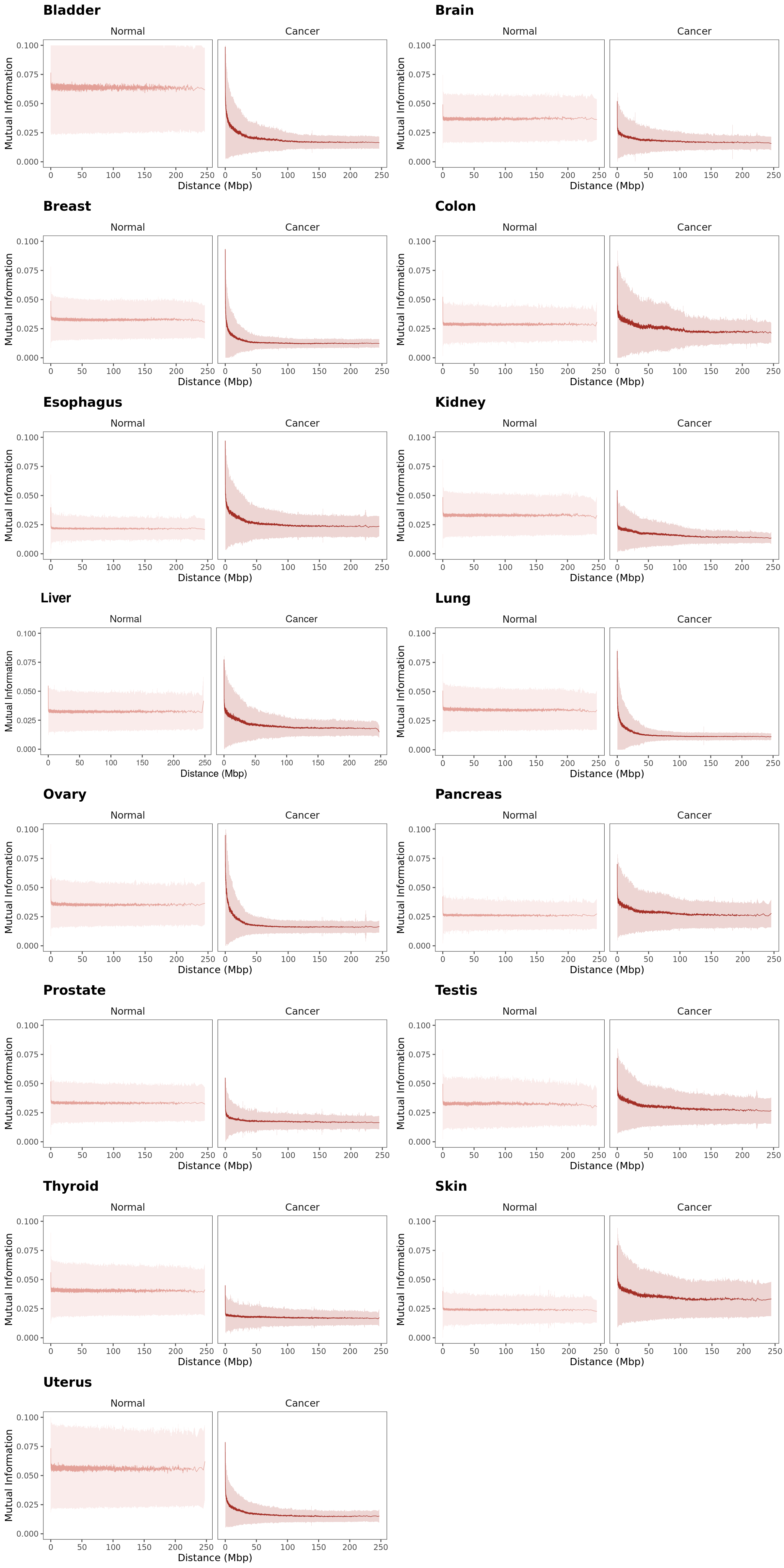

### Supplementary Figure 3

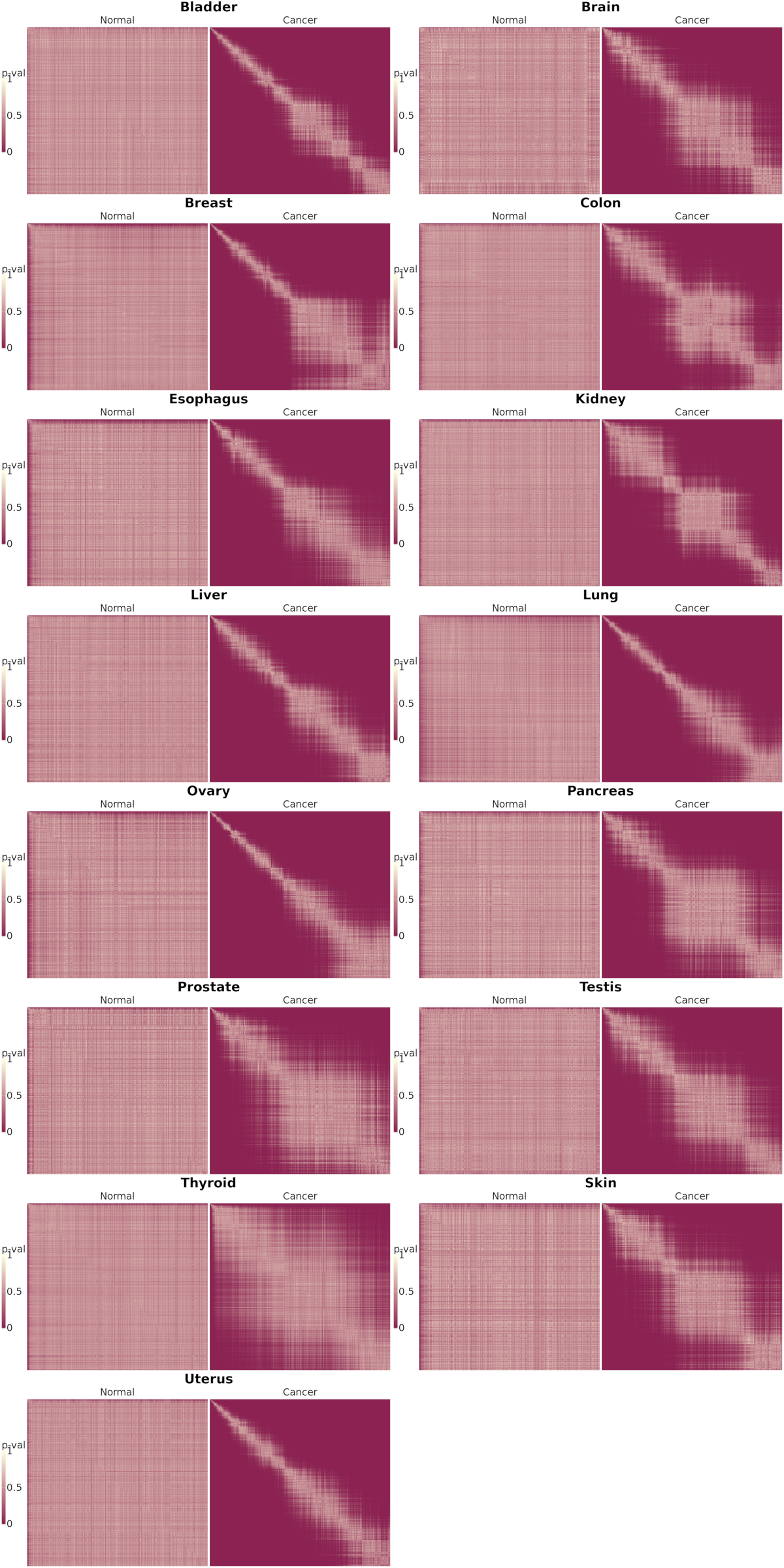

### Supplementary Figure 4

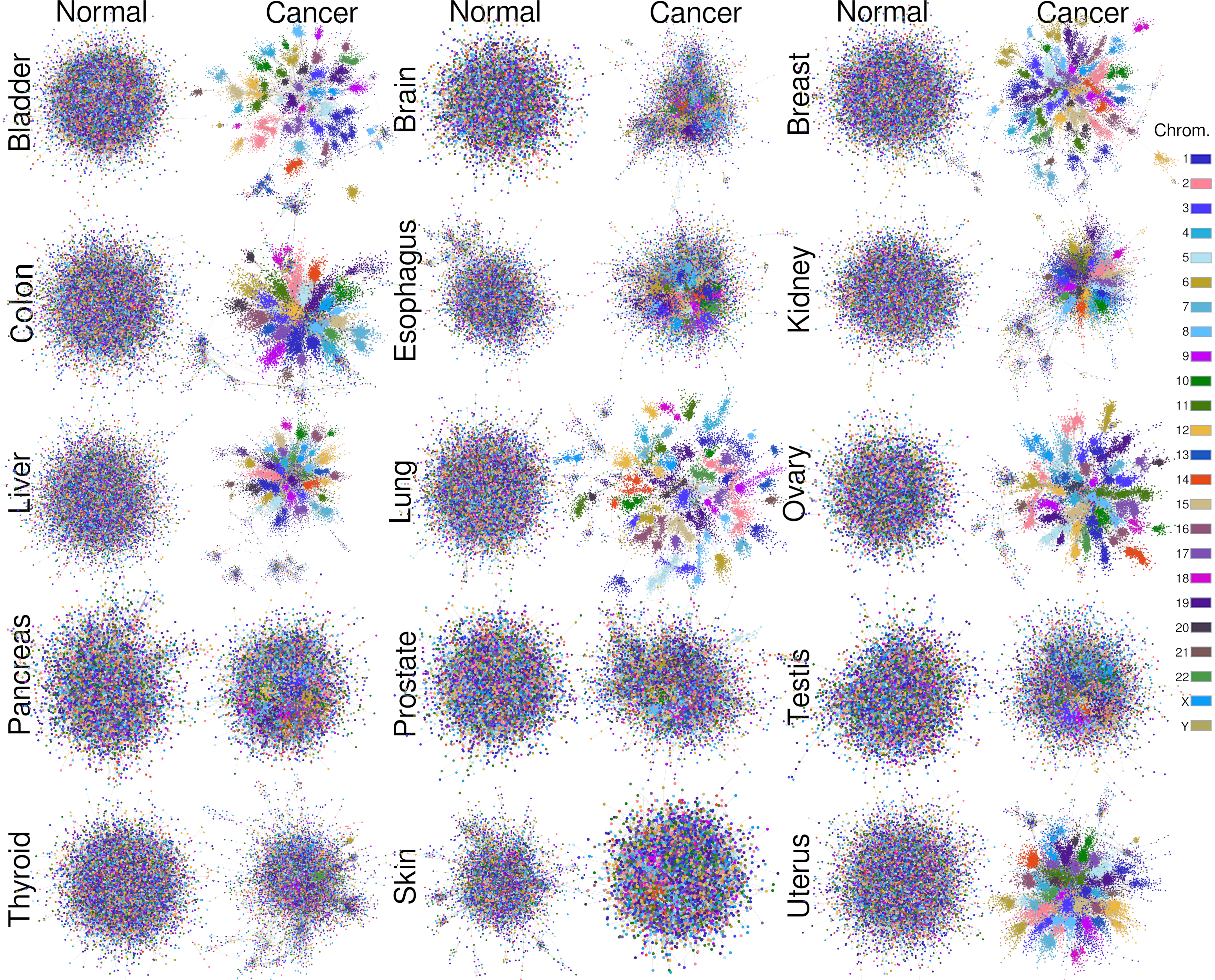
